# Dynamics of mound repair behavior in termites

**DOI:** 10.1101/2025.05.19.654859

**Authors:** Sreekrishna Varma Raja, Amritansh Vats, Parmeshwar Prasad, Chandan Kumar Pandey, Sanjay. P. Sane

## Abstract

Termites of the species *Odontotermes obesus* are found widely across the Indian peninsula. Their presence is visible due to the large over-ground mounds that they build across the landscape. These mounds house their entire colony, including major and minor workers, soldiers, the queen and king, and their brood. In addition, the termites also farm a specific variety of fungus within the mounds, which helps them break down the cellulose-rich food. The precise function of the mound architecture is the subject of some debate, and it has been argued that its structure serves physiological functions including thermoregulation or gas exchange for the entire colony. We hypothesized that if the mound structure indeed serves an important function, termites must be able to sense and repair any injury to their mound structure. To test this hypothesis, we created a hole in the mound wall and observed the dynamics of ensuing breach repair. The breach is quickly detected and repaired by the termites along a consistently sigmodial pattern involving the exponential recruitment of termites during the initial phases of repair, and a derecruitment of termites in the final phases. This behavior provides us with an assay to investigate various phenomena pertaining to mound-building, and to examine the sensory cues involved in detection and repair of the mound. From the basic considerations about the stimuli involved, we developed a logistic model to describe breach repair behavior, highlighting the critical role of a systemic cue in initiating this response. Through a series of experiments utilizing various behavioral assays, we identified the presence of light as this systemic cue that triggers mound repair. When light enters the mound, it triggers construction activity in termites, even though the major and minor workers responsible for repairs do not have image-forming eyes. This suggests that termites detect light through an extraocular mechanism

## Introduction

The ability to control their living environment provides strong adaptive benefits to organisms. To achieve this, many animals including birds (e.g. Collias and Collias, 2014), reptiles (e.g. Hénaut and Charruau, 2012), rodents (Jirkof, 2014), primates (Fruth, Tagg, and Stewart 2018), fish (Rushbrook, Dingemanse, and Barber 2008) and diverse insects (e.g. Sane, Ramaswamy, and Raja 2020) use surrounding materials or self-secreted substances to build nests within which they rear their offspring (e.g. Hansell and Hansell 2005; Hansell and Ruxton 2008). When their nests are damaged, these animals may often repair rather than rebuild them. However, rebuilding a damaged nest requires identifying the exact site of damage to initiate repairs, and recognizing when the breach has been fixed. In eusocial animals that build collectively, such repair processes also requires nest mates to communicate with one another. For these reasons, the study of nest building and repair offers a compelling system in which to study the emergent properties of collective behaviour. These in turn provide inspiration for robotic applications seeking to develop cooperative robots (Rubenstein et al. 2014; Petersen et al. 2012).

Mound-building termites stand out as some of nature’s most remarkable architects and have captivated human interest for centuries (Deshmukh 2001; Marais 2009). Termites, the eusocial members of the class Blattodea, are among the rare non-Hymenopteran examples of eusociality (Emerson 1938; Krishna et al. 2013a; Smeathman 1781). Although the majority of termites live in subterranean nests, many species build large over-ground mounds inside which they house their entire colony. Mound-building termites are present on almost all continents. In India, the termite *Odontotermes obesus* (Kushwaha 1956) occurs across peninsular India, and builds closed mounds reaching heights of up to 3 metres above ground (Krishna et al. 2013b). Despite their small size (∼3 mm), these termites build their impressive mounds through coordinated collective activity. Termites spend the majority of their life within these mounds or subterranean foraging tunnels, emerging only when building the mound or foraging for wood. Similar to other mound building termites, *O. obesus* farms within the mound a fungal garden, which is their primary source of nutrition. Except for the alate termites which take nuptial flights and mate to generate new colonies, all the other termite castes lack image forming eyes.

*O. obsesus* mounds have a rounded tower-like structure, sometimes flanked with fluted side structures (Figure 1A). The function of these mounds remains an open question. Because the termites house their entire colony including their brood within the mound, they must control the internal temperature, air quality and moisture conditions irrespective of changes in the external environmental conditions for the proper growth and maintenance of the colony. Previous researchers have argued that the mound structure is essential for maintaining a constant, uniform internal environment (Emerson 1956; Turner 2009). To achieve this, it is essential for termites to maintain the structural integrity of their mound and to repair any damage to ensure protection from predators and weather. Specifically, termites need to precisely identify the breach location, and rapidly repair it through coordinated, collective action. How do these termites, which lack image-forming eyes, achieve this complex task?

**Figure 1:**
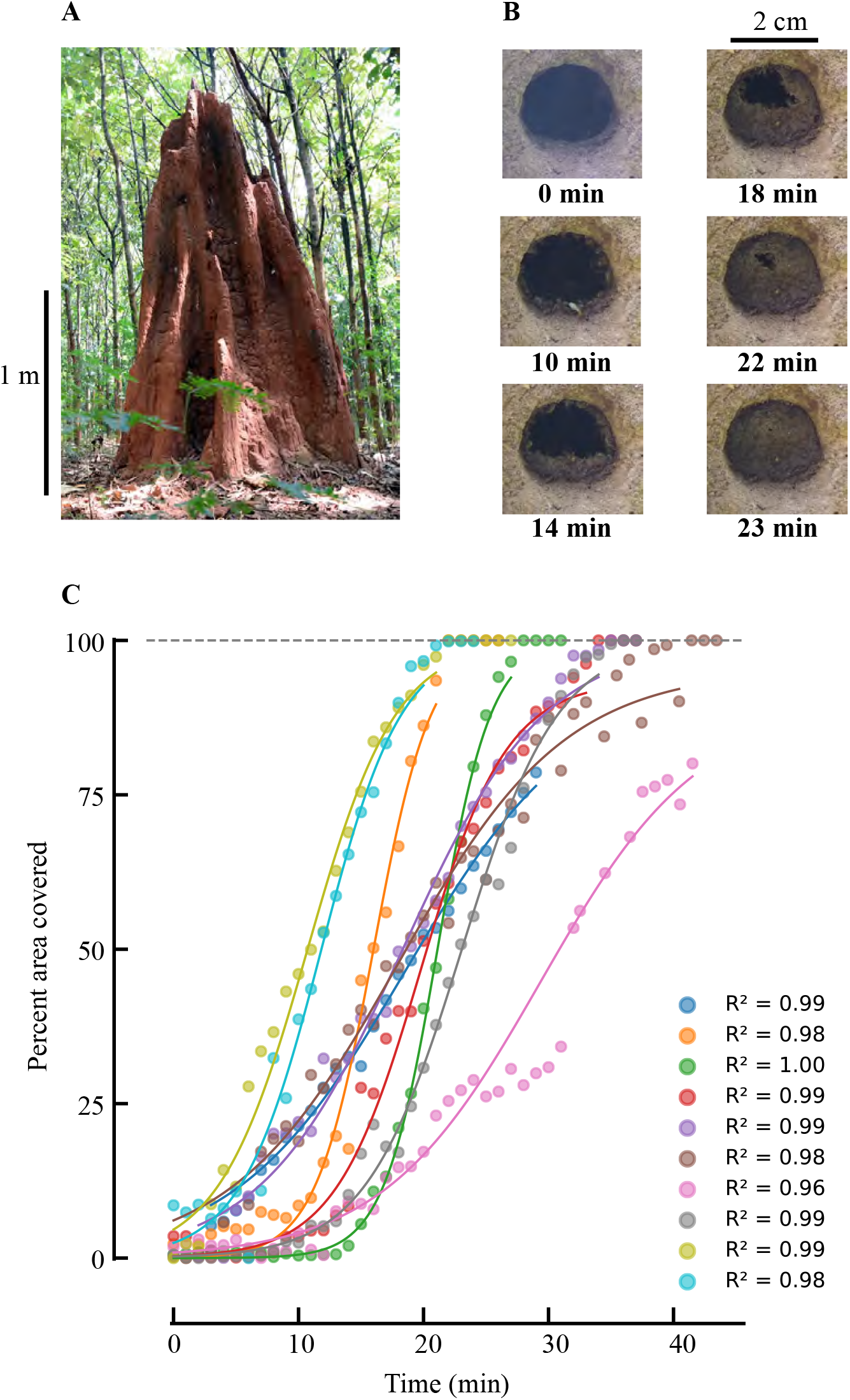
Termite collective construction behavior and quantitative analysis of breach repair dynamics. A. Representative field photograph of a termite mound constructed by *Odontotermes obesus* in its natural forest habitat, showing the characteristic columnar architecture typical of this species. B. Time-lapse sequence documenting the progressive repair of a standardized 2-cm circular breach in the mound surface. Images were captured at 0, 10, 14, 18, 22, and 23 minutes after breach creation, demonstrating the coordinated filling behavior of worker termites. C. Quantitative analysis of repair dynamics showing percentage of breach area covered as a function of time across multiple experimental trials (n=10). Each colored curve represents data from an independent trial fitted to a sigmoidal growth model. The high coefficient of determination values (R^2^ = 0.96-1.00) indicate that the repair process follows predictable sigmoid kinetics. Data points represent actual measurements while continuous lines show the best-fit models.

Here, we show that mound-building termites can sense and repair their mounds when the mound structure is physically damaged (Grassé 1959; Smeathman 1781; Stuart 1967). We developed a series of experimental assays to monitor termite behaviour in response to mound damage, and to understand the underlying mechanisms of repair. With inputs from these assays, we developed a mathematical model in which a logistic equation captures the process by which termites are recruited and de-recruited sigmoidally to the repair site. In this model, we hypothesized that a systemic cue initiates mound building behaviour, which is further sustained by communication between termites. We show that despite lacking image forming eyes, termites are capable of sensing the light entering their mound, which may act as the systemic cue to signal the presence of a breach in the mound wall. Together, these data show that termites have evolved robust mechanisms to collectively maintain their mounds, and also that they possess extra-ocular light sensing abilities.

## Results

To monitor the initiation and time course of breach repair in termites, we made a circular hole in the mound wall (diameter 2 cm, also see Methods). When termites identified this breach, they initiated a repair process in which major and minor worker termites were recruited to the site of damage. Collectively, these termites closed the breach in time frames ranging from 15-30 minutes (Figure 1B). We video-recorded breach-repair behavior and extracted the percentage of the breach area rebuilt as a function of time, *a*(*t*)/*a*_0_ in individual trials (*N* = 11), where *a*(*t*) is the rebuilt area and *a*_0_ is the area of the initial breach. In every case the repair trajectory displayed a sigmoidal relationship including an initial lag phase, an accelerating growth phase, and an asymptotic saturation (Figure 1C). To quantify these dynamics, we fitted each trial’s time course to the three-parameter logistic form

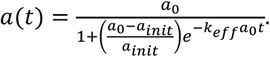

Based on these observations, we hypothesized that the hole in the mound wall presents a systemic stimulus *S*, which is identified by the termites as indicative of a breach. Termites then arrive at the breach site and begin to repair it. Because the number of termites working at the site of the breach were not visible as they are behind the wall, we needed to determine if the dynamics of hole repair reflects progressive recruitment, or whether a fixed number of locally available termites conduct hole repair. If termites work at a steady rate, then the dynamics of the breach repair is indicative of the number of termites.

### The extent of construction is directly proportional to the number of termites

Does the sigmoidal pattern of the breach reflect a similarly sigmoidal recruitment of termites to the repair site, rather than a change in the individual work rate of each termite? Addressing this question required a separate assay in which the termites were visible and could be individually counted as they repaired a breach. Hence, we devised *a cone assay* in which the circular breach was capped with an open funnel, which forced the termites to build on the inner surface of the cone (Figure 2 A, B; see methods). As they deposited soil on the inner surface of the cone, termites could be counted directly from photographic images that were obtained over time while concurrently quantifying the area that they covered with mud (see methods). These measurements indicated a strongly linear relationship (R^2^ values of 11 iterations from lowest to highest: 0.3076, 0.5487, 0.5825, 0.6441, 0.8512, 0.9063, 0.9120, 0.9273, 0.9301, 0.9548, 0.9708) between the number of termites and the extent of soil coverage (Figure 2D). Thus, the extent of construction is directly proportional to the number of termites. This linear relationship also implies that, on an average, each termite maintains a constant work rate. (Figure 2C, E).

**Figure 2:**
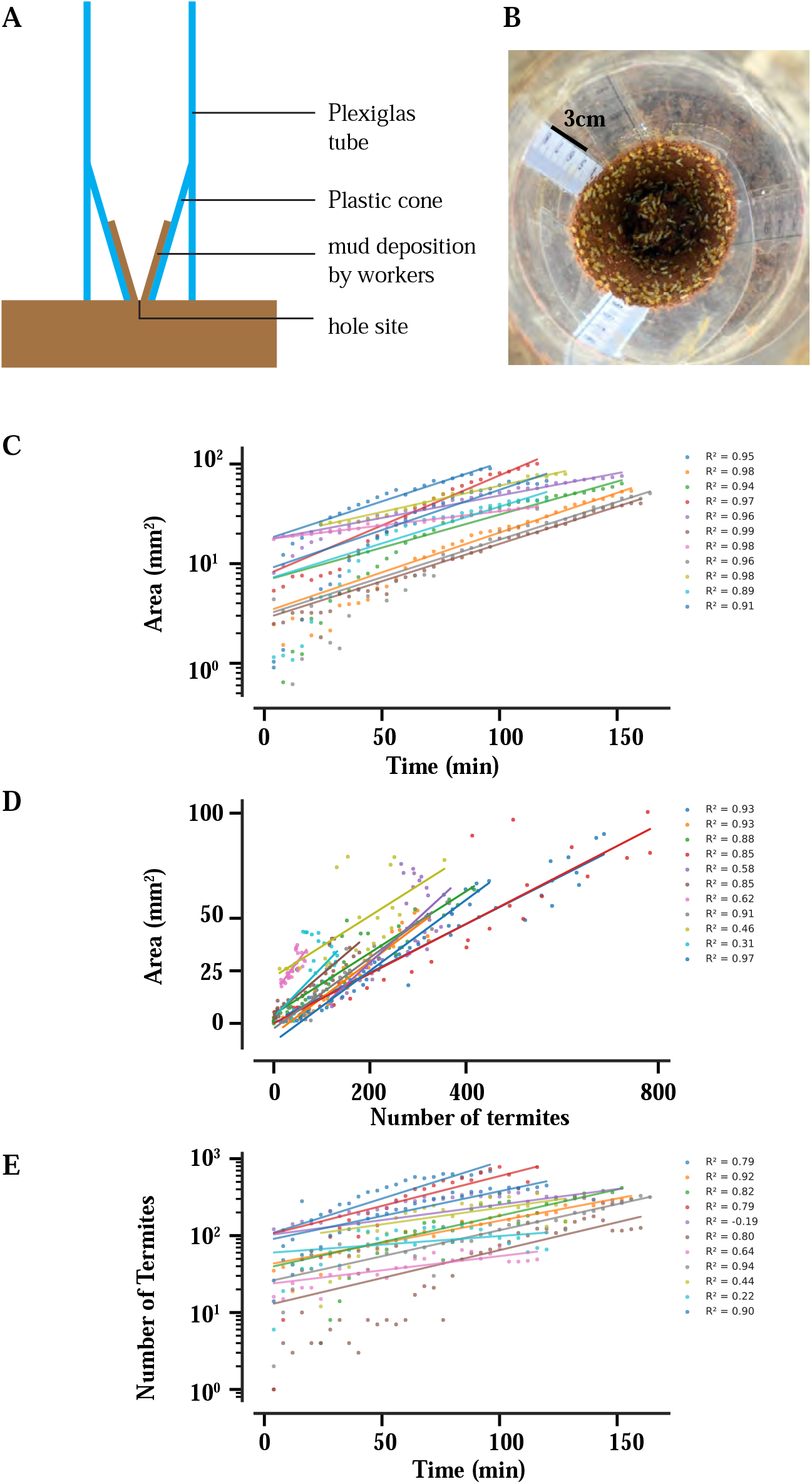
Experimental design and quantitative analysis of termite construction dynamics. A. Schematic illustration of the experimental setup. B. Top-view photograph of active construction, showing termites depositing mud on the cone surface. C. Exponential relationship between mud-covered area (mm^2^) and time (min) across multiple experimental iterations (n=11). Each color represents a different experimental iteration with corresponding R^2^ values shown in the legend. Y-axis is presented on a logarithmic scale. D. Linear relationship between mud-covered area (mm^2^) and number of termites engaged in construction across all iterations. Linear regression fits demonstrate strong positive correlation between worker numbers and constructed area. E. Exponential growth in the number of termites participating in construction over time (min). Y-axis is presented on a logarithmic scale, showing recruitment dynamics across all experimental iterations. The color-coding is consistent across all graphs, with each color representing the same experimental iteration.

### A logistic model captures the dynamics of breach repair in termites

The two experiments described above allowed us to formulate a mathematical model for breach repair which is outlined below.

Let *a*(*t*) = area filled at time *t, n*(*t*) = number of termites present at time *t, a*_*0*_ = size of the hole at time t=0. First, as demonstrated by the cone assay (Figure 2D, E), termites stay and build at a constant rate after arriving at the breach site. This means that the amount of built-up surface indicates the number of termites present at the breach site. If *λ* > *0* be a constant of proportionality then,

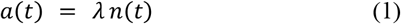

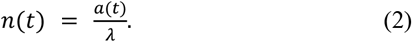

We assume that the creation of a breach causes a systemic *stimulus S*(*t*) such light, temperature gradient, or another cue, which attracts termites from inside the mound to the breach site. After arriving, each termite lays a cue R (e.g. pheromone) which recruits major and minor worker termites from within the mound to the site of repair. As the number of termites *n(t)* increases, the magnitude of R also increases. *Thus, the rate of increase in the number of termites through recruitment is directly proportional to the number of termites present at the breach site and the stimulus strength*.

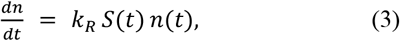

where *k*_*R*_ > *0* is a *recruitment coefficient*.

As the hole is sealed, the effective stimulus also decreases. For simplicity, let the rate of change in *S* be proportional to the rate at which new termites arrive,

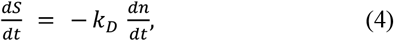

where the *derecruitment coefficient k*_*D*_ > *0*, such that each newly recruited termite reduces the external cue. Integrating, we obtain

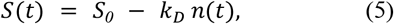

where *S*_*0*_ is chosen so that *S*(*0*) = *S*_*0*_. Hence,

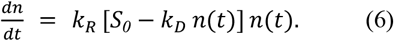

which may be rewritten as

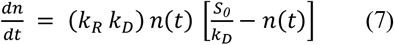

Thus *n*(*t*) follows a *logistic* form with a carrying capacity 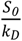.

Because 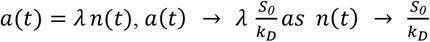.

Physically *a*(*t*) → *a*_*0*_. Thus

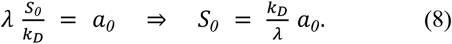

Substituting 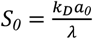 into 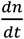:

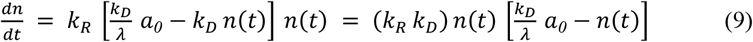

*Thus n(t) has the logistic solution with maximum* 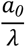. *Finally, since a*(*t*) = *λ n*(*t*),

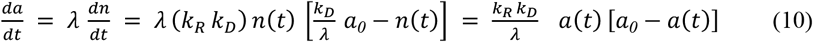

Hence *a*(*t*) follows a *logistic* equation:

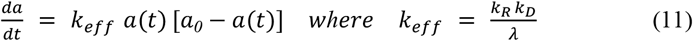

To solve this logistic differential equation, we integrate by separating variables, assuming *a*_*0*_ ≠ *0*:

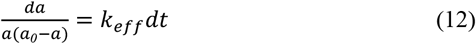

Using partial fraction decomposition:

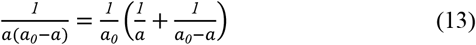

Integrating both sides yields:

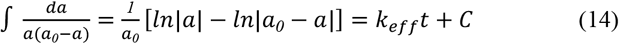

where C is the integration constant. This can be rewritten as:

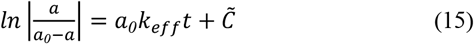

where 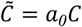. Taking exponentials and assuming *a*(*t*) remains between 0 and *a*_*0*_ (allowing us to drop absolute value signs):

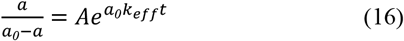

where 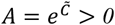. Solving for *a*(*t*):

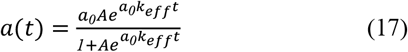

To determine the integration constant A, we apply the initial condition *a*(*0*) = *a*_*init*_:

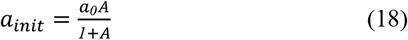

which gives:

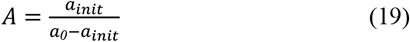

Substituting back yields the classical logistic growth form:

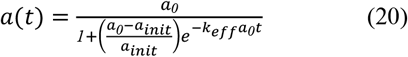

This solution describes how the repaired area *a*(*t*) grows sigmoidally from its initial value *a*_*init*_ and asymptotically approaches the maximum area *a*_*0*_. When *a*(*t*) is small, the growth rate *da*/*dt* remains low because few termites are present. Similarly, when *a*(*t*) approaches *a*_*0*_, the rate becomes small as little unrepaired area remains. The parameter *k*_*eff*_ = *k*_*R*_*k*_*D*_/*λ* governs the speed of repair, incorporating the recruitment rate (*k*_*R*_), derecruitment due to decreasing stimulus (*k*_*D*_), and the proportionality between termite numbers and construction rate (*λ*).

To account for the delay before termites reach the breach site, we modify the solution to include an activation time *t*_*int*_:

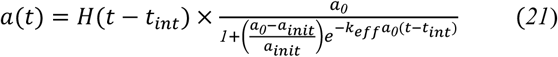

where *H*(*t* − *t*_*int*_) is the Heaviside step function:

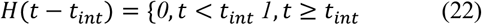

This formulation captures the observation that repair begins only after the first termites discover and respond to the breach, introducing the characteristic lag phase observed in our experimental data.

The above model assumes that the systemic cue S signals termites to arrive at the breach site. Because the light levels inside the mound are extremely low, the presence of light signals the presence of a breach in the mound wall. We therefore tested the hypothesis that light may constitute the systemic cue S that stimulates termite arrival at the breach site. Note that both worker and soldier termites lack compound, image forming eyes. Thus, any sensitivity to light suggests an extra-ocular mode of light sensing.

### Breach repair behaviour is light dependent

To test if termites respond to light levels during breach repair, we devised a transparent tube assay (see methods) in which involved attaching a hollow transparent tube at the mouth of the breach site. This caused termites to enter the tube and build within it, following their instinct to cover foreign surfaces in mud. We created two breaches at equal height (Figure 3A), and covered one breach with a transparent tube, whereas the other was covered with a tube covered in black paper and hence opaque to light. The tubes were left attached to the breach for 24 hrs. If termites are sensitive to light, we predicted that they would build differently in transparent vs opaque tube.

**Figure 3:**
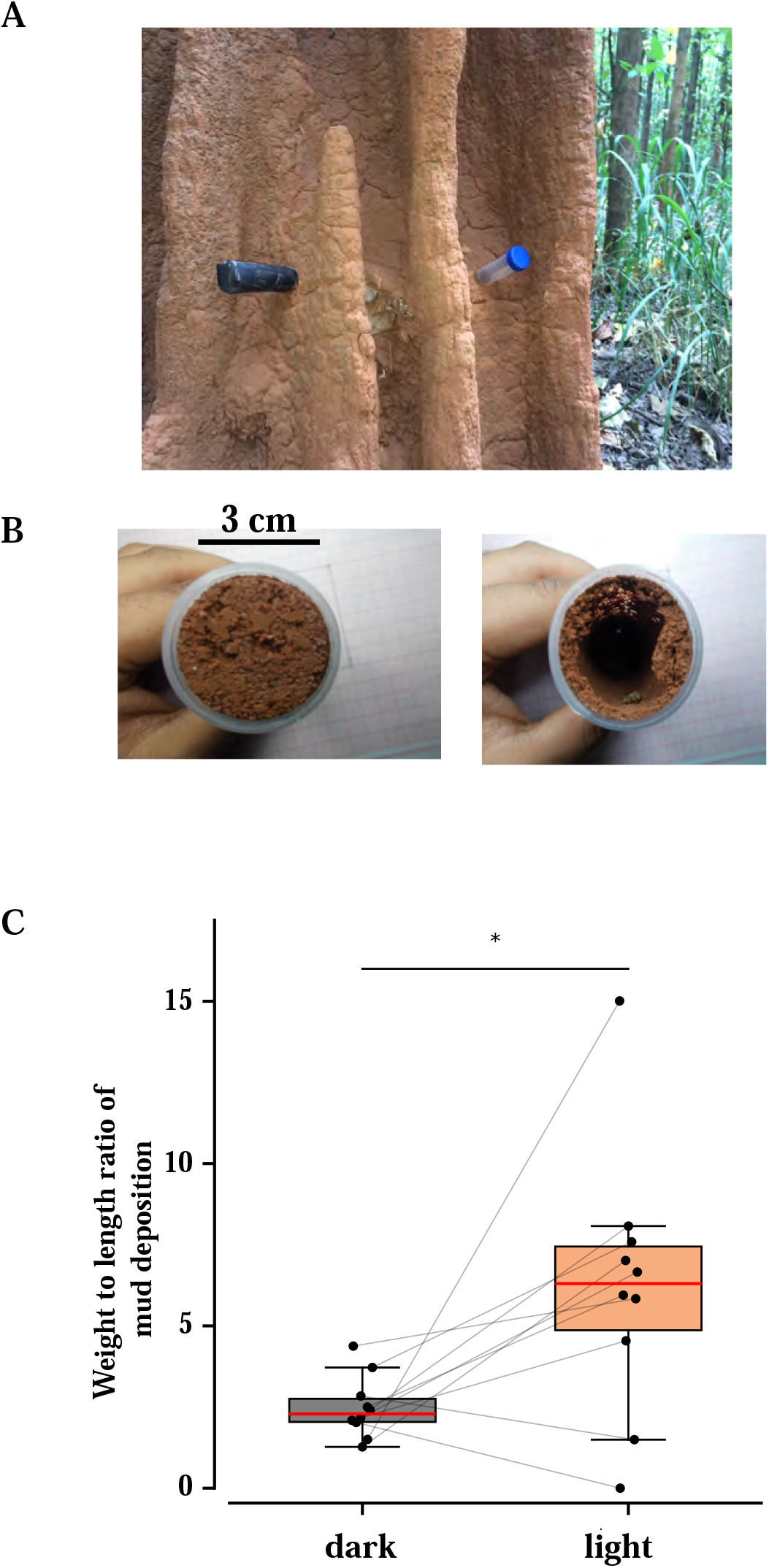
Differential termite construction behavior in response to light exposure. A. Field setup showing two identical plastic tubes inserted into a termite mound at equivalent heights from the base. The tubes differ only in light permeability: one opaque (dark, left) and one transparent (light, right). Both tubes were left undisturbed for 24 hours. B. Cross-sectional view of construction patterns within the tubes after the 24-hour experimental period. Left: The opaque tube shows characteristic hollow-centered construction pattern. Left: The transparent tube exhibits solid mud filling. C. Quantitative comparison of weight-to-length ratio between constructions in dark (gray) and light (orange) conditions. Boxplots display the median (red line), interquartile range (box), and range (whiskers) of measurements. Individual data points (n=10 per condition) are shown as black circles.

In 08/10 trials, we observed that the opaque tube was lined along the walls with mud while the insides remained hollow (Figure 3Bi), whereas the transparent tube was fully packed with mud (Figure 3Bii). The weight of mud per unit length was significantly greater in light-exposed tubes (mean ± SD = 6.22 ± 4.05 g/cm) than in dark tubes (mean ± SD = 2.49 ± 0.95 g/cm); p = 0.024, paired t-test, mean difference (light-dark) = 3.73. Thus, although lacking in image forming eyes, termites built differently within a transparent tube as compared to an opaque tube.

In the above assay, we could not rule out slight differences in temperatures within the two tubes. Hence, to further test this hypothesis, we devised a parallel plate assay in which we measured building rates in two faces of a double parallel plate set up. One face was exposed to bright lights and the other was exposed to extremely low light levels. A conducting aluminium plate that separated the two faces, ensured that the temperature on both sides was the same (Figure 4A). We introduced 30 termites in both sides and measured their mud deposition over a 3 hour duration. The images were photographed and analysed using ImageJ (Figure 4B). This was plotted for different experimental trials for comparison. Although termites showed some building in both faces, the side facing light (normalised to 1.00) was significantly more built up than the side that faced darkness (0.846 ± 0.111) ; p = 0.001, paired t-test, mean difference (light-dark) = 53.86 mm^2^. (Figure 4C).

**Figure 4:**
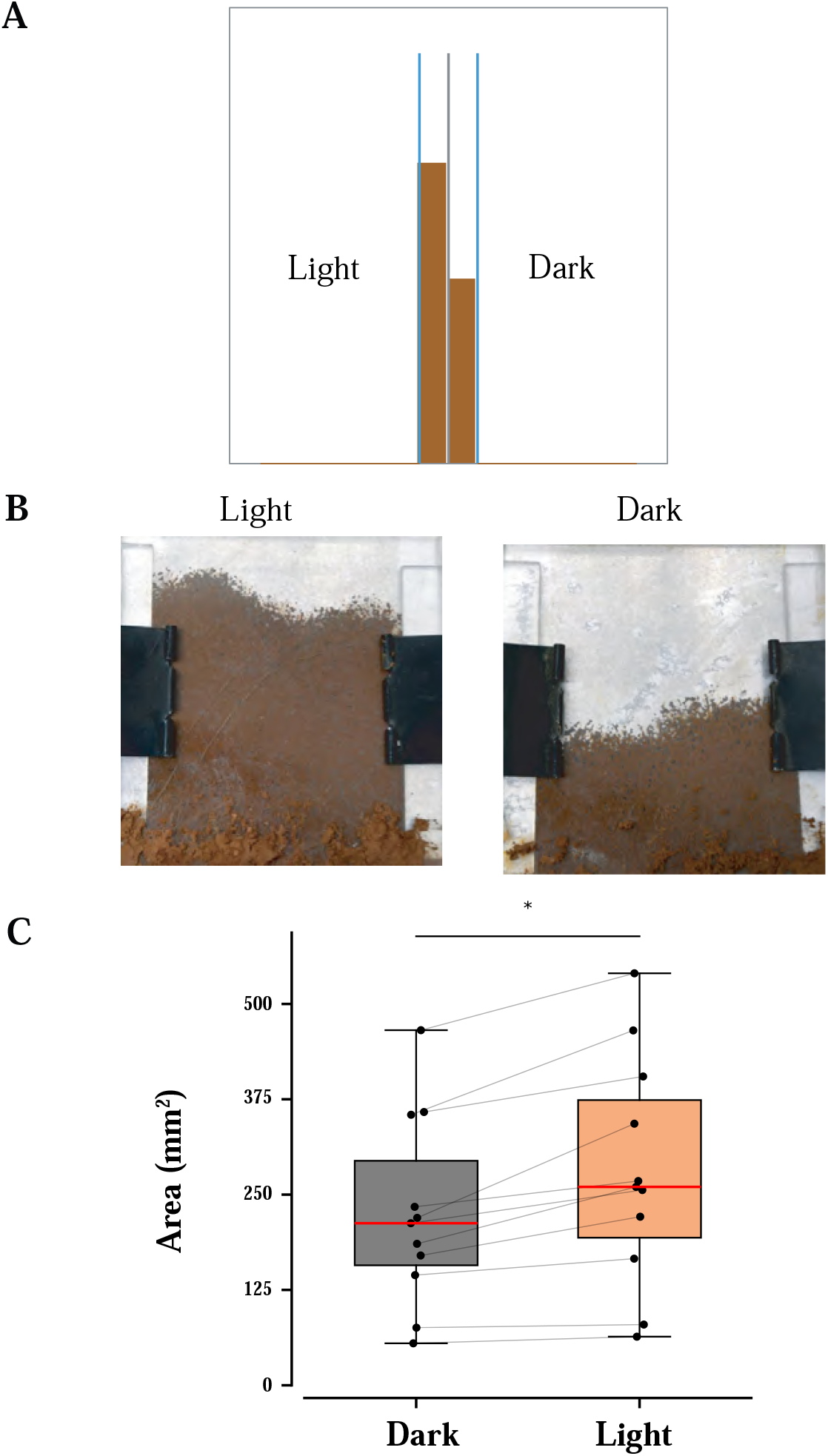
Experimental setup and analysis of termite construction behavior under contrasting light conditions. A. Schematic diagram of the experimental apparatus: a double-spacer parallel-plate arena with an aluminum plate sandwiched between two transparent plates, designed to create controlled light exposure conditions for termite construction experiments. B. Representative photographs showing differential termite construction under light and dark conditions C. Quantitative comparison of construction area under dark and light conditions. Box plots display the median (red line), interquartile range (boxes), and range (whiskers) of measured areas. Individual experimental replicates are shown as black dots (n = 11 per condition).

In the above two assays, the only difference between the two setups was the amount of light in the building area. In both cases, the termites consistently built more in light than in dark, suggesting that the presence of light is a crucial cue which alters the building behaviour in termites, as compared to termites in the dark. Thus, light is a key systemic cue that elicits mound repair response in termites.

## Discussion

In the behaviour investigated here, the presence of a breach in the mound wall elicits a repair behaviour that requires the coordinated efforts of multiple nestmates within the termite mound. Termites need to first detect the presence of a breach, which is followed by recruitment of other nestmates to the breach site and a repair behaviour which eventually closes the breach, which in turn causes the repair behaviour to cease. Because these termites lack image forming compound eyes, it has not been clear how they detect the presence of the breach.

The data presented here offer two novel insights. First, that the dynamics of hole repair follows a logistic equation, involving both the recruitment and derecruitment of termites at the site of the breach. This means that, in addition to the systemic cue that enables termites to identify a breach, the termites have a system of communication that enables them to communicate with their nest mates and recruit them to the breach site. The exact nature of this communication system remains to be described, but it most likely involves pheromone cues that are laid down by each termite as they arrive at the breach site. This explains the initial exponential increase in termite numbers at the breach site. Second, our experiments with transparent tube assay and double parallel plate assay demonstrate that termites respond to light cues even though they lack image forming compound eyes. This means that they possess an extraocular mechanism to detect light. Although extraocular light sensing has been described in many insects, its presence in mound-building termites offers unique insights into how these termites, which normally reside in total darkness, perceive and respond to light in their natural world. Thus, the termites display a very elegant system of detecting and repairing breaches in their mound.

In construction activities by social insects, coordination plays a crucial role, as the overall architecture emerges from the collective efforts of colony individuals. This construction can span multiple generations or involve individuals separated by significant distances. Self-healing, the capacity of a system to discover, diagnose, and respond to disruptions in a cooperative distributed agent environment is exemplified in the nest repair behavior by social insects. While Dawkins’ concept of extended phenotype (Dawkins 1982; Turner 2004) focuses on how genes extend their influence beyond individual organisms, Wilson’s notion of superorganisms (Emerson 1939; Wilson and Sober 1989) delves into collective behaviors within a biological context. These principles are applicable to swarm robotics (Brambilla et al. 2013; Bayindir 2016) and development of autonomous agents that collaborate and exhibit emergent behaviors. In the realm of architectural dynamics within social insects, the scientific landscape is characterized by a notable asymmetry, with a prevalence of theoretical constructs but relative scarcity of empirical observations.

The construction of termite mounds serves as an example of coordination during construction (Deneubourg and Franks 1995) The role of volatile pheromones released by queen to guide workers in soil deposition is an example of coordinated building (Bruinsma 1979; Bonabeau et al. 1998; Jones 1979; Deneubourg 1977). But do termites coordinate building activity away from the queen nest? Although the phenomenon of mound repair in termites has been known for more than two centuries (Smeathman 1781), a measurement and analysis of the repair dynamics offers surprising insights into the underlying mechanisms.

## Methods

### Field sites

*Odontotermes obesus* are commonly found and widely distributed in parts of South India [4]. For the study described here, we used the termite mounds on the campus of NCBS, Bangalore, India (Figure 1A). All experiments were performed in the mounds on the field. To access termites for specific experiments, we carved out a small hole (3 cm diameter) in the mound skin, which provided a natural mix of termites from different castes. We did not control the number or caste of termites entering the experimental arena.

### Hole repair assay

In this assay, a breach was made in the mound wall by carving out a circular hole of 2 cm diameter using a small garden pick axe. Typically, after a brief period of time, soldier termites arrived and positioned themselves around the breach. Following this, worker termites arrived at the breach and began repairing it by depositing small pellets of mud that were excavated from around the breach site, ingested, processed, regurgitated and rolled using their mandibles (Supplementary movie 1). As the breach in the mound was repaired by the termites, the size of the hole diminished with time (Figure 1B). This repair process was monitored using Sony Handycam (DCR-DVD108, Tokyo) mounted on a tripod directly facing the mound. Using ImageJ software (National Institutes of Health, Bethesda, Maryland, USA), these videos were analysed frame-by-frame using the contrast between the hole and the surrounding mud to identify and measure the hole size as it diminished over time. In repeated trials, we took care to make the breach at different locations because termites are known to substantially remodel parts of their mound that are vulnerable to repeated injury. After measuring both the original breach and the breach remaining after repair, we calculated the ratio of the repaired breach as a function of time (*breach repair ratio*).

### Cone assay

In the hole assay described above, it was possible to quantify how much construction work was accomplished by monitoring the *breach repair ratio*. However, it is not possible to see the termites at work because they were usually behind the wall of the breach and hence not always visible. To establish the correlation between the number of termites and the actual amount of soil deposited by them at the working site, we needed an assay that forced the termites to build in the open, thus enabling us to physically count the number of termites working and measure the deposited mud. To achieve this, we took a small, hollow plastic frustum, inverted it and fixed it inside a longer hollow plexiglass tube. This setup was affixed on top of a hole (2 cm diameter) in the mound skin such that the hole remains contiguous with the mouth of the inverted frustum (Figure 2A). The presence of an extended surface around the hole in a confined environment elicits deposition of mud pellets and building along the inner surface of the frustum, which could then be filmed. We fixed a Nikon 5200 DSLR camera to view the process of building from the top angle which enabled us to see all the termites working (Figure 2B). A time-lapse was set for the camera to acquire pictures every 4 minutes. Using the count tool of Adobe Photoshop CS5, we manually counted the number of termites building on the frustum surface in each frame.

Termites first make a single pellet thick layer of mud on the surface of the frustum, similar to the thickness of the covering over the breach. Thus, measuring the area of the surface covered by mud directly translates to the quantity of mud deposited. Using a set scale along the length of the frustum while recording, we calculated the area built on the frustum surface.

### Parallel plate assay

To measure the varying rates of building activity through the day, we devised an assay to enable us to record the amount of mud deposited with time at different times of the day. Two thin plexiglass sheets (10 cm X10 cm) were placed facing each other, separated by two plexiglass spacers (4 mm thick) on lateral sides. This parallel plate arena apparatus was fixed in place using binder clips. We then placed this parallel plate arena over a small hole in the mound skin similar to the frustum (Figure 3A). Worker termites were stimulated by the presence of this extended surface that was contiguous with the hole. They began depositing mud pellets on the inner surfaces of the two plates, gradually covering this planar area in time (Figure 3B). We recorded the growth in this area of built with time at different times of the day. The measurement was done till the entire planar arena was deposited with mud, hence seizing the deposition process or that the termites had capped off this building process at a certain height. Using ImageJ tools, we obtained the area of mud deposition on the plates of the arena with time. This was plotted for different experimental trials at different times through the day for comparison.

### Transparent tube assay

To test for the effect of light on building activity, we prepared two types of hollow transparent plastic tubes (diameter=3 cm and length=9 cm) from 50 ml Falcon® tubes with their conic tips sawed off. One end was closed with lid, whereas the open end fit snugly into the hole in the mound (diameter= 3 cm). One tube was covered with black paper along its length and one end to ensure that no light entered into the tube. The second tube was not covered and remained transparent, however the lid was closed at one end as in the previous case. Tubes were mounted in the same mound 30 cm apart (Figure 3A) at the same height and left for 24 hrs. After 24 hrs, the tubes were collected and weighed, to measure the extent to which the tube was filled with mud. We photographically documented the extent of filling of each tube (Figure 3B). At the end of the experiment, we measured the mass of the soil per unit length of both dark and transparent cylinders. Because the tube was not always fully filled, we divided this mass by the length of the tube over which the mud was filled to obtain a weight to length ratio (Figure 3C)

### Double parallel plate assay

As a second test for the effect of light on building activity, we designed an assay in which we used two thin plexiglass sheets (10 cm X10 cm) sandwiching an aluminium plate. These three plates were separated by two plexiglass spacers (4 mm thick) which were held in place by binder clips to form a double parallel plate arena. On one side, an aluminium foil was used to prevent light from entering thus keeping that side in darkness. The base of the assay was contiguous with the mound environment. Thirty termites were introduced into both sides and we measured their mud deposition over a 3 hour duration. To analyse the images, we used ImageJ as previously described for the parallel plate assay. This was plotted for different experimental trials for comparison.

### Statistical Analysis

All statistical analyses were performed using Python 3.8 with SciPy (v1.7.1), NumPy (v1.20.3), and statsmodels (v0.12.2) packages. Figures were generated using Matplotlib (v3.4.3). To quantify the sigmoidal repair trajectory, we fitted each experimental trial to a three-parameter logistic model using robust nonlinear least-squares with a Soft L1 loss function to down-weight residual outliers. Parameter constraints were implemented (0≤a_0_,a_init_≤100, 10^−4^≤k_ell_≤10) to ensure biological plausibility. Prior to fitting, we removed all data points at or above 99% saturation to avoid bias from the asymptotic phase. Goodness-of-fit was assessed by the coefficient of determination (R^2^) and the root-mean-square error (RMSE) expressed as a percentage of the fitted carrying capacity. Model parameters were compared across replicates using descriptive statistics (mean ± SD).

Linear regression was used to assess the relationship between the number of termites and the extent of construction in the cone assay. The exponential growth of termite number and area built over time was analysed using semi-log plots with linear regression on log-transformed data. R^2^ values were calculated to determine the strength of these relationships. For the transparent tube assay, differences in mud deposition between light and dark conditions were analysed using paired t-tests (p = 0.001) with a mean difference (light-dark) of 3.73 (Figure 3). For the double parallel plate assay (Figure 4), paired t-tests revealed significant differences between light and dark conditions (p = 0.024) with a mean difference (light-dark) of 53.86 mm^2^. All statistical tests were two-tailed with significance threshold set at α = 0.05. Data are presented as mean ± standard deviation unless otherwise specified.

## Notes

### Competing Interest Statement

The authors have declared no competing interest.

### Funding

- Human Frontier Science Program: GP0066/2012
- Department of Atomic Energy: 12-R&D-TFR-5.04-0800,

